# Transcranial alternating current stimulation attenuates BOLD adaptation and increases functional connectivity

**DOI:** 10.1101/630368

**Authors:** Kohitij Kar, Takuya Ito, Michael Cole, Bart Krekelberg

**Author notes:** McGovern Institute for Brain Research, Department of Brain and Cognitive Sciences, Massachusetts Institute of Technology, Cambridge, MA. K.K., and B.K designed the experiments. K.K. carried out the experiments. K.K., and T.I. performed the data analysis. K.K., M.C. and B.K. wrote the manuscript. The authors declare no competing financial interests. All correspondence should be addressed to Bart Krekelberg.

## Abstract

Transcranial alternating current stimulation (tACS) is used as a non-invasive tool for cognitive enhancement and clinical applications. The physiological effects of tACS, however, are complex and poorly understood (Liu et al. 2018). Most studies of tACS focus on its ability to entrain brain oscillations (Herrmann et al. 2013), but our behavioral results in humans (Kar and Krekelberg 2014a) and extracellular recordings in nonhuman primates (Kar et al. 2017) support the view that tACS at 10 Hz additionally affects brain function by reducing sensory adaptation. Our primary goal here was to test this hypothesis using BOLD imaging in human subjects. Using a motion adaptation paradigm developed to quantify BOLD adaptation (Huk et al. 2001) and concurrent fMRI and tACS, we found that tACS significantly attenuated adaptation in the human motion area (hMT+). In addition, an exploratory analysis showed that tACS increased functional connectivity between the stimulated hMT+ and the rest of the brain, in particular the dorsal attention network. We conclude that weak 10 Hz currents applied to the scalp affect both local and global measures of brain activity.

**New and Noteworthy:** Concurrent transcranial alternating current stimulation (tACS) and fMRI show that tACS affects the human brain by attenuating adaptation and increasing functional connectivity. This work is important for our basic understanding of what tACS does, but also for therapeutic applications, which need insight into the full range of ways in which tACS affects the brain.

## INTRODUCTION

Weak alternating currents applied to the scalp modulate behavior (Antal et al. 2008; Helfrich et al. 2014a; Kar and Krekelberg 2014b), but the mechanistic route from currents on the scalp via changes in neural activity to behavioral change is far from understood (Liu et al. 2018). Our goal is to develop insight into the neural level changes caused by transcranial alternating currents, and ultimately use this insight to improve the transcranial stimulation technique. Current experimental evidence and computational models support two modes of action. The first is entrainment: alternating currents can entrain ongoing oscillations (Francis et al. 2003; Fröhlich and McCormick 2010; Ozen et al. 2010; Zaehle et al. 2010; Ali et al. 2013; Krause et al. 2019) and, given sufficiently long stimulation periods, this entrainment can outlast stimulation (Helfrich et al. 2014b; Kar 2015; Strüber et al. 2015). At the cellular level, entrainment is thought to result from the sub-threshold modulation of the membrane potential by the weak intracranial electric field generated by the applied currents (Herrmann et al. 2013). Previous studies combining BOLD imaging with tACS have focused primarily on this mode of action. The experimental results, however, are not equivocal, with some reporting decreased BOLD signals in task-active areas (Vosskuhl et al. 2016) others increases in BOLD signals in task inactive areas (Cabral-Calderin et al. 2016b), or decreases in BOLD after tACS offset (Alekseichuk et al. 2016). Recently, we identified a second mode of action; extracellular recordings in macaque middle temporal cortex (MT) showed that tACS at 10 Hz attenuates spike frequency adaptation (Kar et al. 2017). This requires only a brief period of stimulation (3 s), and primarily affects neurons that are actively responding to the sensory input. These two properties potentially enhance the temporal and functional specificity of this mode of action. Based on in-vitro recordings (Fernandez et al. 2011), we have speculated that this mode of action relies on tACS-induced membrane voltage fluctuations and the consequent inactivation of Na^+^ or Ca^2+^ dependent K^+^ channels (Kar et al. 2017). Our specific goal in the current study was to bridge the gap between animal and human studies and to provide evidence of this second mode of action – the attenuation of adaptation – in the human brain. We pursued this question by quantifying BOLD signal changes evoked by sensory (visual motion) adaptation in the presence or absence of tACS. Previous fMRI studies have identified an area in the human brain (hMT+) that, just as MT in the macaque, is highly selective for visual motion (Tootell et al. 1995). The BOLD response of this area is typically reduced after prolonged exposure to moving patterns (Huk et al. 2001), and this so-called BOLD adaptation is generally accepted as a reflection of neuronal adaptation (Krekelberg et al. 2006a), similar to what is observed in single neurons in the macaque (Krekelberg et al. 2006b; Kar and Krekelberg 2016). Our experiments confirmed the prediction that tACS attenuates BOLD adaptation and thereby provide direct support for tACS’ second mode of action in the human brain. Beyond this immediate goal of providing support for our specific adaptation hypothesis, we also used our whole-brain data set to demonstrate that tACS increases functional connectivity. These analyses add to the growing evidence that weak currents applied to the scalp can affect brain activity in complex ways. Overall, we argue that although tACS clearly does modulate brain activity much more work is needed to understand its properties and thereby develop a technique that may be able to target specific cortical areas, identified networks, or brain functions.

## RESULT

Based on prior behavioral findings in humans (Kar and Krekelberg 2014b) and electrophysiological data in nonhuman primates (Kar et al. 2017), we have argued that tACS at 10 Hz attenuates sensory adaptation. Here we tested this hypothesis by measuring the influence of tACS (±0.5mA, 10 Hz) on adaptation of the BOLD response in hMT+.

### tACS reduces adaptation in hMT+

We adopted the method of Huk et al (Huk et al. 2001) to measure the strength of direction-selective adaptation in hMT+ (Figure 1). Subjects first view one direction of motion (the adaptation direction; outward drifting gratings) for several seconds. The relatively well-documented properties of neuronal adaptation (Krekelberg et al. 2006a; Kar and Krekelberg 2016) predict that the BOLD response to a subsequently presented pattern with the same direction of motion (outward) is smaller than the BOLD response to the opposite, non-adapted direction of motion (inward drifting gratings). We alternated blocks of trials in which patterns moved in the adapted or non-adapted direction, and quantified direction-selective BOLD adaptation using the correlation between the predicted alternation, and the observed BOLD response (DS; Methods). Consistent with the findings of (Huk et al. 2001), our bilateral adaptation stimulus led to significant BOLD adaptation in hMT+ of both hemispheres (mean Pearson r(i.e. DS) ± SD; left : DS=0.48±0.018, right: DS=0.48±0.017; t-test; t(9) = 39; p<0.001). To test our hypothesis that 10 Hz tACS reduces adaptation, we placed one stimulation electrode approximately over left hMT+ (PO7 in the 10-20 system) and the other over the vertex (Cz). This montage matched our previous behavioral study (Kar and Krekelberg 2014b). Based on individualized head models (Methods), the average electric field magnitude in a sphere centered on left hMT+, was 0.17 ± 0.03 V/m (mean ± standard deviation across subjects), while right hMT+ received 0.09 ± 0.01 V/m. The magnitude of these estimated fields depends strongly on the (coarse) estimates of tissue conductivities used in the model, and should therefore be taken with a grain of salt. The ratio of the field strength in the left vs. right hMT+ is more robust against errors in these assumptions. Therefore, we prefer the quantification in terms of a ratio: left hMT received a field that was 1.9 ± 0.4 times stronger than the field in right hMT+ (range: 1.2 to 2.7). For convenience, we refer to the left as the stimulated hemisphere and the right the non-stimulated hemisphere. In these terms, our experimental prediction was that stimulation applied during the presentation of the adapting stimulus (the outward moving grating) should reduce adaptation more in the stimulated hMT+ than in the non-stimulated hMT+.

**Fig. 1.**
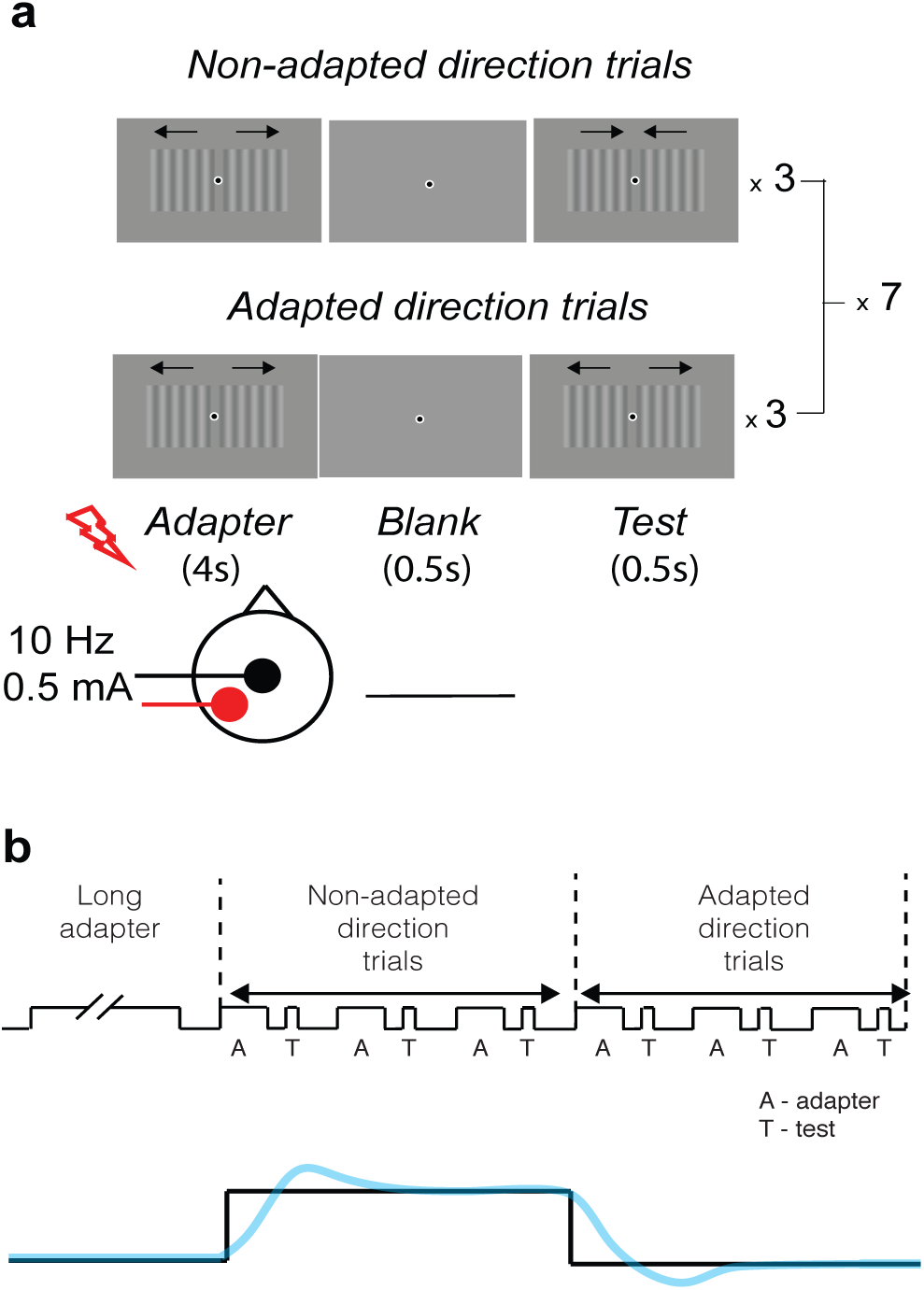
Experimental paradigm. Subjects fixated on the central dot throughout. a) Adaptation was induced with outward drifting gratings, these were presented first for 30 s at the start of a block (long adapter), and then for 4 s (top-up) in each trial. The adapted responses were measured with gratings moving in the same (adapted direction) or opposite (non-adapted) direction. A set of three non-adapted direction trials was alternated with sets of three adapted direction trials (always with the same adapter), and this set of 6 trials was repeated 7 times in one block. b) Schematic of the predicted neural and BOLD time course. Because adaptation typically reduces the neuronal response (Krekelberg et al. 2006a; Kar and Krekelberg 2016), the response in adapted-direction trials is predicted to be smaller than that in non-adapted direction trials. At the block level, adaptation should therefore result in a higher neural activity in the non-adapted direction trials than in the adapted-direction trials; the prediction for the expected BOLD signal (blue) follows from the assumptions underlying the hemodynamic response (Methods).

Figure 2a shows the strength of adaptation (DS) from one example subject in the tACS ON trials. The red clusters show the voxels that adapted significantly (mean Pearson r(169) = 0.1; p<0.001). Clearly, there were fewer significantly adapted voxels in the stimulated left hemisphere than the non-stimulated right hemisphere. To assess the statistical significance of changes in adaptation strength, we used a 4-way mixed effects model with fixed factors time, tACS, and hemisphere, and subject as a random factor, and focused solely on the interaction effects to control for nonspecific changes with time (Methods). The significant interaction between tACS and hemisphere (F(1,1) = 5.01; p=0.0001) confirms that adaptation was indeed weaker in the stimulated than the non-stimulated hemisphere. To visualize this, Figure 2b shows a direct comparison of the adaptation strength in each of the hemispheres in the tACS ON condition.

**Fig. 2.**
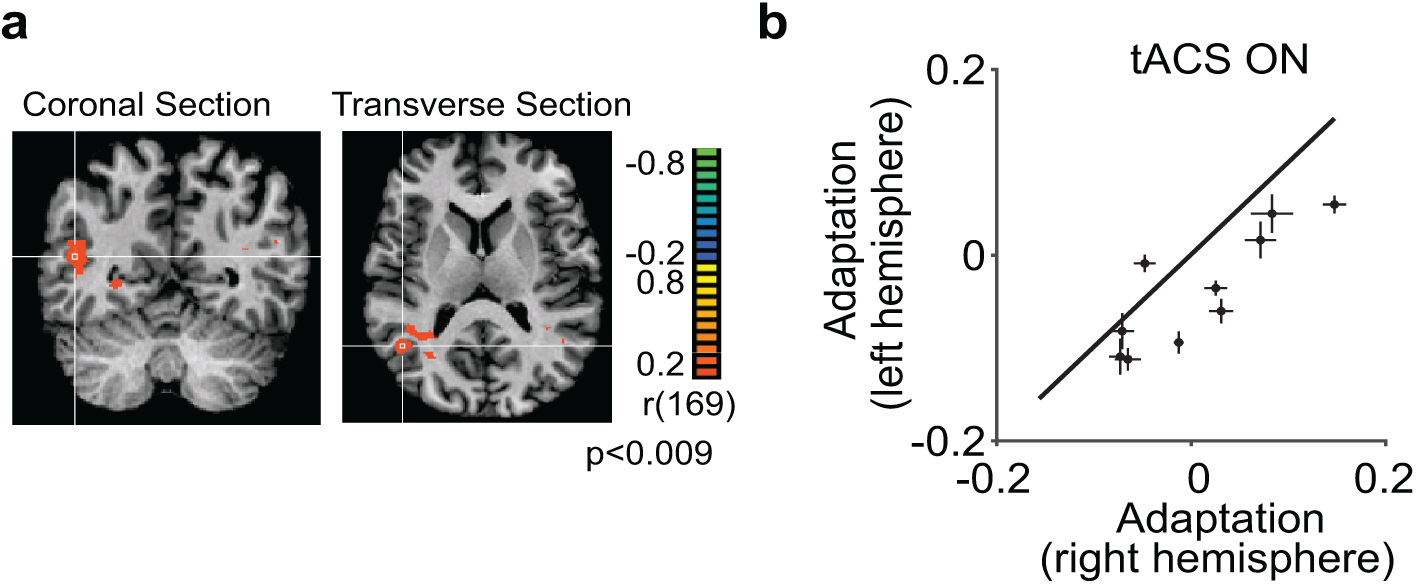
tACS reduces BOLD adaptation. a) Results from an individual subject. Color maps show the amount of BOLD adaptation (DS) as quantified by the correlation between the BOLD signal and the adaptation predictor (see Methods). Images are in neurological convention. The white cross hairs in the coronal and the sagittal slices are at Talairach coordinates x = 40, y= −72, z = −7. hMT+ in the non-stimulated (right) hemisphere shows significant adaptation, but adaptation was non-significant in the stimulated (left) hemisphere. b) The average BOLD adaptation (DS) in the stimulated hemisphere (y-axis) as a function of the BOLD adaptation in the non-stimulated hemisphere (x-axis), separately for each subject. Error bars show bootstrapped 95% confidence intervals.

One mechanism for the attenuation of adaptation could be that 10 Hz tACS reduced neural (spiking) activity (Vosskuhl et al. 2016) and thereby indirectly reduced adaptation because neurons that respond less, typically adapt less (Kar and Krekelberg 2016). This explanation predicts that BOLD signals should be lower during tACS. We evaluated this by comparing the BOLD response in hMT+ during the long adapter stimulus in the trials when tACS was on, with those in which tACS was off. A two way repeated measures ANOVA with hemisphere (stimulated/non-stimulated) and tACS (ON/OFF) as factors showed that tACS was associated with a increase, not a decrease, in BOLD response (main effect of tACS: F(1,9) = 7.8, p = 0.02). This finding is incompatible with a model in which reduced neuronal firing during tACS results in reduced adaptation (measured after tACS offset). Instead, we speculate that tACS interferes directly with the cellular mechanisms underlying adaptation (Kar et al. 2017).

### Functional Connectivity

Although the primary goal of our experiments was to test the adaptation hypothesis, the whole-brain data allowed us to perform an exploratory analysis of the influence of tACS on functional connectivity. Because our task was designed to drive visual motion areas, we investigated only the extent to which tACS changed the functional connectivity of hMT+ (Methods). At the whole-brain level, we computed weighted degree centrality (WDC) (also known as global brain connectivity (Cole et al. 2010, 2012; Ito et al. 2017)) for the stimulated hMT+. This graph theoretic measure represents the average FC of a region to the entire brain [29]. tACS at 10 Hz significantly increased the WDC of hMT+ (F(1,9) = 13.89; p = 0.005). At the network level, we computed the average FC of hMT+ to a set of pre-defined functional networks (Power et al. 2011). FC increased significantly between hMT+ and the dorsal attention network (DAN) (F(1,9) = 16.20; p = 0.04; FDR-corrected; Fig. 4b,d), but not (13.90>F; p>0.05; FDR-corrected) with any of the other 12 functional networks defined by (Power et al. 2011). We also asked whether this increase in FC could be attributed statistically to specific areas, but none of the individual regions of interest defined by (Power et al. 2011) increased FC significantly after multiple comparison corrections (21.65>F; p>0.05; FDR-corrected).

**Fig. 3.**
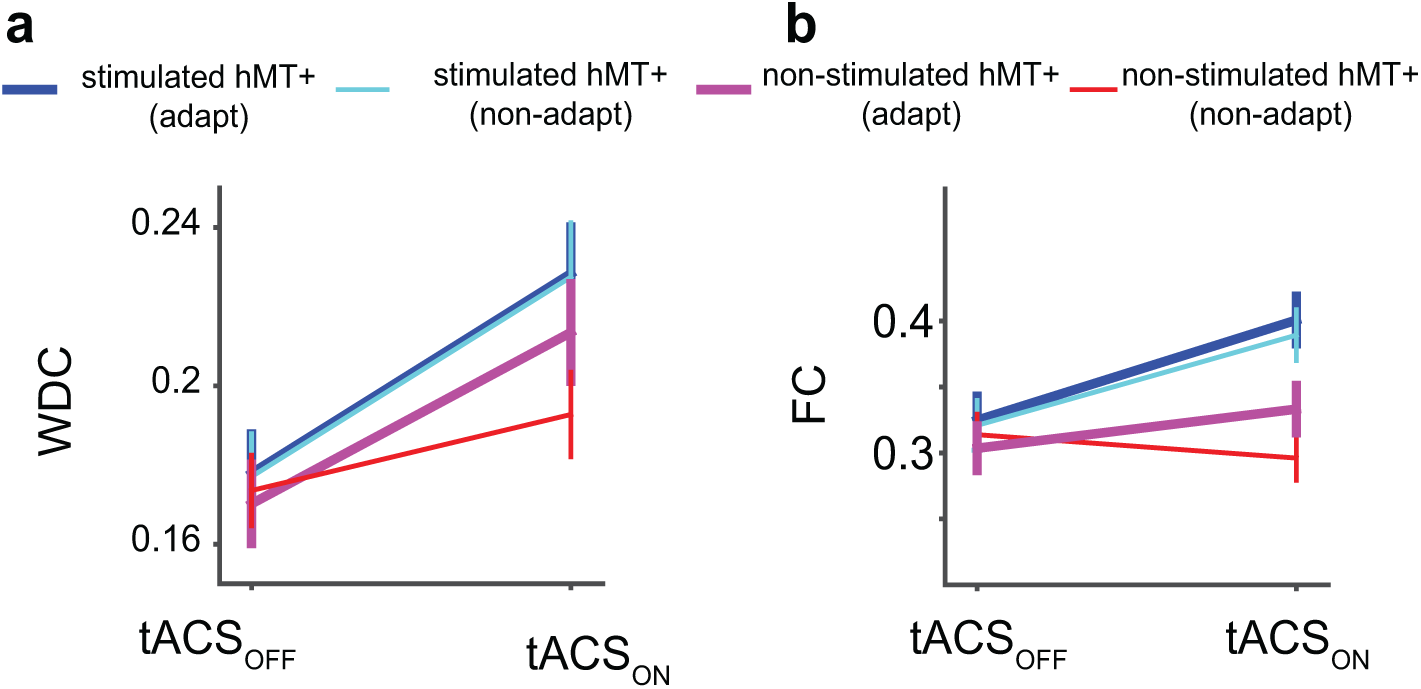
tACS increases functional connectivity. (a) Weighted degree centrality of hMT+ increased with tACS; more so in the stimulated hemisphere than the unstimulated hemisphere, and, at least in the stimulated hemisphere, there was no interaction with adaptation. (b) Functional connectivity between hMT+ and the dorsal attention network increased with tACS, more in the stimulated than the non-stimulated hemisphere. Adaptation did not affect this interaction. These analyses show that 10 Hz tACS increases functional connectivity, and that at least some of this effect (b) does not depend on adaptation.

**Fig. 4.**
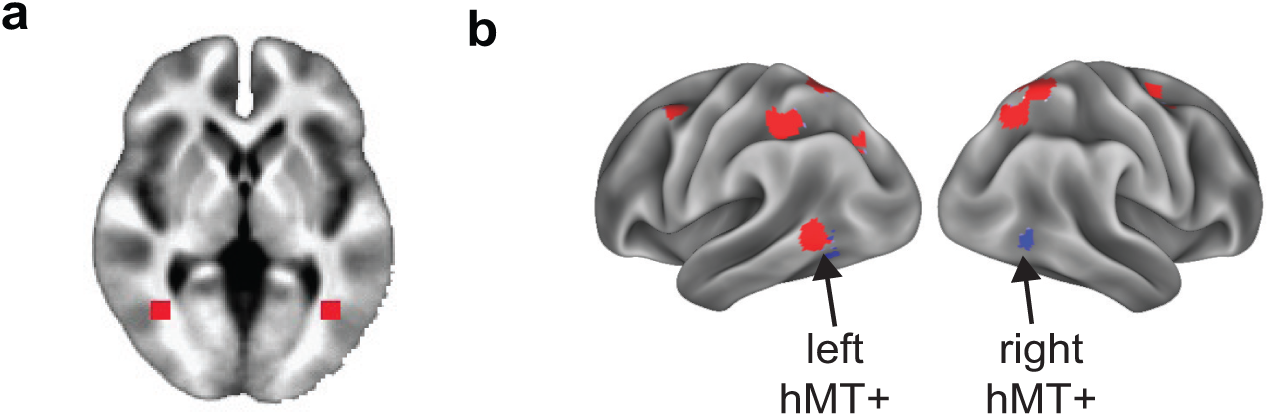
Functional connectivity analysis. (a) Bilateral hMT+ seeds in volume space with a 5mm radius at (40,-60,0) and (−40,-60,0). (b) Surface visualization of bilateral hMT+ seeds (blue) and DAN regions (red). Average FC from the hMTI+ ROIs to the DAN network increased significantly after tACS (p=0.005).

## Discussion

We investigated how tACS affects BOLD signals in area hMT+ during the processing of a visual motion stimulus. Consistent with our predictions based on behavioral and electrophysiological data, we found that tACS reduced adaptation. We also observed that the application of tACS increased functional connectivity between the hMT+ and the rest of the brain, and the dorsal attention network in particular. We first address some of the potential confounding factors and limitations in the interpretation of our data and link our findings to previous studies using concurrent fMRI and tACS. Then we speculate on the neural mechanisms that could be responsible for the tACS-effects we reported, and conclude with a brief discussion of the implications of our findings for the future use and interpretation of tACS-effects.

### Confounding factors

#### Phosphenes

Application of tACS produces phosphenes via retinal stimulation(Schutter and Hortensius 2010; Kar and Krekelberg 2012; Laakso and Hirata 2013). Phosphenes can act as a distractor and thereby reduce attention, which reduces adaptation (Rezec et al. 2004). However, this generalized effect would apply to both hemispheres. The fact that attenuation of adaptation in the left (stimulated) hemisphere was larger than in the right (non-stimulated) hemisphere therefore controls for this confound and we conclude that the attenuation of adaptation is not a side effect of phosphenes, or any stimulation-induced overall changes in arousal or attention.

#### Artifacts introduced by tACS in the scanner

Transcranially applied electric fields in the MRI scanner can produce artifacts in the EPI signal on the scalp and in the cerebrospinal fluid (Antal et al. 2014). Those artifacts, however, occur during direct current stimulation, not during (40 Hz) alternating current stimulation. This suggests that the BOLD signals we measured are unlikely to be affected greatly by stimulation artifacts. In addition, we applied tACS both during adapted direction trials and non-adapted direction trials. Because artifacts would be generated in both sets of trials, they do not contribute to the differences between these trials that underlie our main result (the reduction in adaptation).

#### Targeted Stimulation

Large stimulation electrodes are appealing because they result in low current densities, which limits tactile and/or painful sensations on the scalp. However, the electric field calculations show that such electrodes generate fields that spread widely across the scalp and in the brain (Opitz et al. 2016; Huang et al. 2017; Alekseichuk et al. 2018). In our specific montage, the fields in the target area (left hMT+) were twice as large as those in the opposite hemisphere. In the macaque (Kar et al. 2017), we used an analogous stimulation approach, but measured the fields intracranially. There the target area received an electric field that was four times larger than that in the opposite hemisphere. Given the numerous technical differences between these studies and the species differences, this seems a reasonable level of agreement. Both studies show that some spatial targeting can be achieved even with large pad electrodes, but off-target stimulation that is 25-50% of the targeted stimulation magnitude can be expected even in the opposite hemisphere. Interestingly, both our electrophysiological and our fMRI data provide evidence that these two- to four-fold differences in field strength, even at a low (< 1 V/m) overall magnitude, are sufficient to induce measurable differences in neural activity. This upside (small field differences result in measureable activity differences) also has a downside in that it limits the interpretation of TCS experiments. For instance, we cannot infer (based on these data alone) that the stimulation of hMT+ directly causes the reduction in behavioral measures of adaptation (Kar and Krekelberg 2014b), because off-target stimulation (i.e. fields outside left hMT+) is not negligible compared to on-target stimulation. Such restrictions on the interpretation of causality will apply to most, if not all, TCS experiments. Notably, even if stimulation targeted to brain area X results in a behavioral change, that does not prove that X is causally involved in the behavior.

#### tACS mechanism

These results provide novel support for our hypothesis that tACS attenuates adaptation. First, they provide the first evidence that tACS reduces neural adaptation in the human brain as it does in the nonhuman primate (Kar et al. 2017). Second, using BOLD imaging we could analyze the effects of tACS during stimulation, a period we could not consider in the nonhuman primate recordings due to the electrical artifacts in the sensitive electrophysiological recording hardware (Liu et al. 2018). The finding that the BOLD signal increased rather than decreased during stimulation supports the view that tACS interferes with the induction of adaptation, rather than neural activity per sé. The current data do not address the mechanistic details at the cellular level, but we’ve previously speculated that small membrane voltage fluctuations may interact with the dynamics of the Na+ and Ca2+ activated K+ channels that underlie visual adaptation, analogous to findings in the hippocampus (Fernandez et al. 2011). Testing this hypothesis likely requires in-vitro recordings, or the use of transgenic animals in which these channels can be expressed selectively (Stroud et al. 2012). Our finding that tACS increased the BOLD response to the onset of a visual stimulus appears to conflict with previous reports demonstrating a decrease in stimulus-driven BOLD response (Cabral-Calderin et al. 2016a; Vosskuhl et al. 2016). One interesting potential explanation of this discrepancy is that our montage primarily targeted parietal cortex, while earlier studies Oz/Cz montage targeted occipital cortex. Occipital cortex is dominated by alpha oscillations, and their power correlates negatively with the BOLD signal (Scheeringa et al. 2012). This suggests that tACS entrainment of alpha could reduce the BOLD signal, but only in early visual cortex (Vosskuhl et al. 2016). Such areal specificity of tACS’ modes of action are intriguing and potentially powerful as they suggest that cortical targeting could be achieved not just by choosing appropriate electrode montages, but also by the selection of particular stimulation frequencies (Cabral-Calderin et al. 2016a).

#### Functional Connectivity

We showed that apart from attenuating adaptation, 10 Hz tACS also strengthened the functional connectivity of hMT+. The spatial specificity of this effect (primarily in the DAN) suggests that this is not solely due to the spatial spread of the tACS fields (which would predict increased FC primarily near the stimulation electrodes). Alternatively, the entrainment of an intrinsic 10 Hz oscillation in hMT+ may have led to the entrainment of other areas via normal neuronal communication pathways. This of course raises the question why the DAN was specifically affected, and whether other stimulation frequencies might have led to changes in FC between different areas. Because we investigated FC within the context of a very specific motion task, we limited ourselves to the analysis of hMT+ seeds, and our data cannot answer such questions. Recent work using resting-state FC, however, has cast a much wider net and found that tACS-induced changes in FC depend in a highly complex fashion on brain areas, stimulation frequencies, and electrode montages (Cabral-Calderin et al. 2016b). Unravelling this complexity is necessary to further the goal of using tACS to modulate brain networks.

## Conclusion

Our results show that tACS applied during prolonged visual motion stimulation increases activity in hMT+ and yet reduces the influence of the prolonged exposure on subsequent responses. We conclude from this that tACS attenuates the induction of adaptation in the human brain as it does in the monkey brain (Kar et al. 2017). In addition, we found an increase in functional connectivity of hMT+ with the rest of the brain, and the dorsal attention network in particular. Our data show that these modes of action are at least partially non-overlapping. We speculate that making an area less adapted, more active, and more strongly connected with the attention network, could all contribute to the cognitive enhancements reported using tACS.

## Materials and Methods

### Subjects

Ten subjects (5 female) participated in the study. Subjects gave written consent and all had normal or corrected to normal vision. This study was conducted according to the principles expressed in the Declaration of Helsinki and approved by the Institutional Review Board of Rutgers University.

### tACS

We combined transcranial current stimulation with MRI acquisition, which has previously been shown to be safe, and results in minimal artifacts and loss of signal to noise (Antal et al. 2014). The stimulus generator was in the control room and was connected to the electrodes on the subject’s head via wall-mounted radio frequency filters, and MR compatible, shielded cables (MRIRFIF and custom CBL200, Biopac). The electrode leads were equipped with a 5.6 kOhm resistor to limit RF heating of the head. In addition, we placed each lead in a neoprene covering to avoid overlapping wires and wire loops, and thus limit current induction. The leads were passed out through the side of the head coil and then led along the bore towards the back of the scanner. We applied tACS using an STG4002 stimulus generator (Multi Channel Systems, Reutlingen, Germany). The circular stimulating electrodes (BML Basic Physician’s Supply, Inc) were made of conductive rubber (7.6 cm diameter), and attached to the scalp using Signa electrode gel (Parker Laboratories Inc., Fairfield, NJ, USA). One electrode was placed at the canonical location of left hMT+ (PO7-PO3 in the 10-20 system), the other on the vertex (Cz). The current intensity amplitude was 0.5 mA (i.e. 1 mA peak-to-peak); the frequency was 10 Hz. These parameters were chosen to match our previous behavioral experiments in humans (Kar and Krekelberg 2014b) and electrophysiological recordings in the macaque (Kar et al. 2017).

### Apparatus

A Canon REALiS SX80 Mark II LCOS projector (60 Hz) back-projected the stimuli onto a screen located at the end of the MRI bore. Subjects viewed the stimuli via a mirror attached to the head coil. The combined distance of the screen to the mirror and the mirror to the subjects’ eyes was 103 cm. The display measured 22° (width) by 12° (height) and had a resolution of 1920 × 1080 pixels. Stimulus presentation and the triggering of stimulation were under the control of in-house, OpenGL-based software.

### Motion adaptation paradigm

We adopted the visual motion adaptation paradigm from Huk et al (Huk et al. 2001) to quantify direction-selective motion adaptation in the BOLD signal. Subjects fixated a dot at the center of the screen while viewing two moving gratings on either side of the dot (5° × 5° centered on + 7°). For each experimental run, both gratings initially moved inward for 30 s (long adapter). Subsequent trials were classified into two conditions. In ‘adapted direction trials’, a top-up adapter (both gratings moving inwards for 4s) was followed by a test stimulus moving inward for 0.5 s. In ‘non-adapted direction trials’, the same top-up adapter was followed by a test stimulus moving outward for 0.5 s. The sequence of trials (i.e., after the initial 30 s long adaptation) alternated between three non-adapted direction trials and three adapted direction trials. Six of these trials were considered one block. Each block was presented 7 times per run. Each subject participated in at least four experimental runs in the same session; two without tACS (tACS-off), followed by two with tACS (tACS-on). In the tACS-on conditions, the current was applied whenever the adapter stimulus (either the long adapter or the top-up adapter) was on the screen (i.e., during the induction of adaptation).

### fMRI

#### Data Acquisition

We conducted all imaging at the Rutgers University Brain Imaging Center (RUBIC) using a 3T MRI (Tim Trio, Siemens) scanner, and a 32-channel head coil with ample padding around the head to minimize head movement. We used a T1-weighted MPRAGE sequence to collect 1 mm3 resolution anatomical images from each subject. For functional scans, we used a T2*-weighted echo planar imaging sequence (repetition time = 2 s, echo time = 25 ms, flip angle = 90°, matrix = 64 × 64). The 35 slices (in plane resolution = 3 × 3 mm; slice thickness = 3 mm) covered the entire brain and were oriented approximately parallel to the line connecting the anterior to the posterior commissure (ACPC).

### Data Analysis

#### ROI Analysis

##### Data Preprocessing

We analyzed the fMRI data with BrainVoyager (version 2.6; Brain Innovation, Maastricht, Netherlands) and MATLAB (MathWorks). We discarded the first nine volumes of each functional scan before preprocessing the functional data. Preprocessing included linear trend removal, slice scan time adjustment, 3-D motion correction with alignment to the first volume within an MRI session, and temporal filtering using a high-pass filter with a 0.0078 Hz cut-off. The functional images were superimposed on the high-resolution anatomical images and incorporated into the 3D data sets through trilinear interpolation. The complete data set was transformed into Talairach space. We defined area hMT+ by a sphere (10 mm radius) around its canonical Talairach coordinates: (40,-60, 0) for the right hemisphere and (−40,-60, 0) for the left hemisphere.

##### BOLD adaptation

Based on the known adaptation properties of MT neurons in the macaque (Krekelberg et al. 2006b; Patterson et al. 2014; Kar and Krekelberg 2016), and previous studies in humans (Tootell et al. 1995; Huk et al. 2001) we predicted that for our choice of stimuli, adaptation would primarily reduce the neural response in the adapted-direction trials compared to the opposite-direction trials. Due to the slow response dynamics of the BOLD signal, this predicts that the BOLD signal should be higher than average in the non-adapted direction trials, and lower than average in the adapted-direction trials. Formally, we computed a predictor (Figure 1b) by convolving the predicted increase in neural activity in the non-adapted direction trials with a two-gamma hemodynamic response function (HRF, onset = 0 s, response to undershoot ratio = 6, time to response peak = 5 s, time to undershoot peak = 15 s, response and undershoot dispersion = 1). We quantified the strength of direction-selective adaptation (DS) for each voxel as the Pearson correlation between this predictor and the BOLD time course of the voxel. In voxels with a positive correlation, adaptation reduced the response; in voxels with a negative correlation, adaptation increased the response.

#### Functional Connectivity Analysis

##### Data Preprocessing

All connectivity preprocessing and analyses were performed using MATLAB and AFNI (version 2011-12-21)(Cox 1996). The first nine volumes of each scan were discarded. EPI images were slice-time corrected, aligned to the subject’s skull-stripped MPRAGE in native space, motion-corrected, and transformed to Talairach space. A linear regression was subsequently performed to remove nuisance parameters from the time series. This included the six motion parameters, and ventricle and white matter time series along with their derivative time series. In addition, to remove any potential spatial co-activation confounds with functional connectivity (FC) analyses, we also regressed out BOLD signals related to stimulus presentation (adapter on/off, test on/off and tACS on/off), all convolved with the same canonical HRF as in the above analysis involving BOLD activity during adaptation. The residual time series was then spatially smoothed within a one-voxel dilated gray matter mask at 6 mm FWHM.

##### ROI-based functional connectivity analysis

Because our paradigm was specifically targeted to drive visual responses (and adaptation) in hMT+, we chose these areas as the seeds for our FC analyses and the 264 pre-defined functional regions of the Power et al (2011) atlas as the target regions. To match the spherical size of the ROIs used in the Power atlas (5 mm radius), we defined area hMT+ by a 5 mm radius sphere around the canonical coordinates at (40, −60, 0) for the right hemisphere and (−40, −60, 0) for the left hemisphere. We removed regions 257 and 262 from the Power atlas for our analyses, since they either overlapped or were adjacent to the left and right hMT+ masks. For each ROI, we first split up the time series according to stimulation condition (tACS OFF and tACS ON) and direction of motion (non-adapted direction and adapted direction). Then, we computed the Pearson correlation from each of the hMT+ to all other regions to obtain FC measures for each of the conditions, resulting in four connectivity vectors for each hMT+. We removed negative FC connections and self-connections, since they likely reflect spurious connections that add noise to the underlying network topology (Rubinov and Sporns 2010). As a first test, we computed the weighted degree centrality for hMT+, a graph-theoretic measure that computes the average FC of a region across the entire brain (Rubinov and Sporns 2010). Next, to obtain more specificity with regards to the functional networks that were driving this whole-brain effect, we computed the average FC from hMT+ to each functional network across the different conditions. Lastly, to obtain region-to-region specificity, we determined FC values from every region in the Power atlas to hMT+.

##### Statistical Analysis

In our experimental design, tACS OFF blocks preceded tACS-ON blocks to prevent after-effects of stimulation from contaminating later BOLD signals. Therefore, we cannot distinguish between a main effect of time and a main effect of stimulation. Given that the passage of time could affect BOLD signals in numerous ways (e.g. subject fatigue, scanner drift) this is a potentially serious confound. For this reason, we followed previous approaches (Cabral-Calderin et al. 2016b) to focus solely on statistical interactions and not main effects. Specifically, for the direction selective adaptation DS, we used a mixed effects model with stimulation condition (tACS ON/OFF) and hemisphere (stimulated/non-stimulated), as fixed effects and subjects as a random-effect. We consider the statistical interaction between hemisphere and stimulation as the effect of interest for direction selective adaptation, because it controls for time and placebo effects of stimulation (e.g. phosphenes and arousal; see Discussion). Similarly, for the FC analyses, we analyzed the correlations using a four-way mixed effects model with direction, stimulation, and hemisphere as fixed effects and subjects as random effects. In this analysis, the three-way interaction between stimulation, hemisphere, and direction indicates a change in FC that could potentially be accounted for by a change in adapted state (i.e. it could be a follow-on effect of the attenuation of adaptation). Therefore, we also searched specifically for a significant two-way interaction between stimulation and hemisphere, but in the absence of the above three-way interaction. All correlation values were Fisher z-transformed prior to statistical analyses. All p-values were corrected for multiple comparisons with the false discovery rate (FDR) procedure.

##### Field Strength

We used the Realistic Volumetric-Approach to Simulate Transcranial Electric Stimulation (ROAST, version 4.11) pipeline (Huang et al. 2019) to model electric field strengths based on the subjects’ individual structural T1-weighted MRI, the linear ICBM average brain (ICBM-152) (all at 1 mm3 resolution) and the ICBM-NY brain (Huang et al. 2016) at 0.5 mm3 resolution. For the individual subjects, all ROAST simulations were performed on their raw T1 images, using tissue conductivity parameters from the literature (Huang et al. 2017) (white matter : 0.126 S/m, gray matter : 0.276 S/m, cerebrospinal fluid: 1.65 S/m, bone: 0.01 S/m, and skin: 0.456 S/m). Once model simulations were complete, we used fMRIPrep (Esteban et al. 2019), version 1.4.1rc1, to determine the spatial transforms necessary to normalize each T1 to the MNI 152 template, and then applied that transform to the modeled electric field magnitude. From these spatially normalized model results, we extracted the electric field magnitude in a sphere with a 10 mm radius, centered on the canonical location of hMT+ in the left and right hemisphere ([±40, −60, −0]).

## ACKNOWLEDGMENTS

We thank Gregg Ferencz and Jasmine Siegel for excellent technical support and Jessica Wright, Melanie Arroyave, Sophia Chirayil, and Heta Patel for assistance during data collection.

## References

Alekseichuk I, Diers K, Paulus W, Antal A. Transcranial electrical stimulation of the occipital cortex during visual perception modifies the magnitude of BOLD activity: A combined tES–fMRI approach. Neuroimage 140: 110–117, 2016.

Alekseichuk I, Mantell K, Shirinpour S, Opitz A. Comparative Modeling of Transcranial Magnetic and Electric Stimulation in Mouse, Monkey, and Human [Online]. bioRxiv. https://www.biorxiv.org/content/early/2018/10/13/442426.

Ali MM, Sellers KK, Fröhlich F. Transcranial alternating current stimulation modulates large-scale cortical network activity by network resonance. J Neurosci 33: 11262–75, 2013.

Antal A, Bikson M, Datta A, Lafon B, Dechent P, Parra LC, Paulus W. Imaging artifacts induced by electrical stimulation during conventional fMRI of the brain. Neuroimage 85: 1040–1047, 2014.

Antal A, Boros K, Poreisz C, Chaieb L, Terney D, Paulus W. Comparatively weak after-effects of transcranial alternating current stimulation (tACS) on cortical excitability in humans. Brain Stimul 1: 97–105, 2008.

Cabral-Calderin Y, Anne Weinrich C, Schmidt-Samoa C, Poland E, Dechent P, Bähr M, Wilke M. Transcranial alternating current stimulation affects the BOLD signal in a frequency and task-dependent manner. Hum Brain Mapp 37: 94–121, 2016a.

Cabral-Calderin Y, Williams KA, Opitz A, Dechent P, Wilke M. Transcranial alternating current stimulation modulates spontaneous low frequency fluctuations as measured with fMRI. Neuroimage 141: 88–107, 2016b.

Cole MW, Pathak S, Schneider W. Identifying the brain’s most globally connected regions. Neuroimage (2010). doi:10.1016/j.neuroimage.2009.11.001.

Cole MW, Yarkoni T, Repovs G, Anticevic A, Braver TS. Global Connectivity of Prefrontal Cortex Predicts Cognitive Control and Intelligence. J Neurosci 32: 8988–8999, 2012.

Cox RW. AFNI: software for analysis and visualization of functional magnetic resonance neuroimages. Comput Biomed Res 29: 162–73, 1996.

Esteban O, Markiewicz CJ, Blair RW, Moodie CA, Isik AI, Erramuzpe A, Kent JD, Goncalves M, DuPre E, Snyder M, Oya H, Ghosh SS, Wright J, Durnez J, Poldrack RA, Gorgolewski KJ. fMRIPrep: a robust preprocessing pipeline for functional MRI. Nat Methods 16: 111–116, 2019.

Fernandez FR, Broicher T, Truong A, White JA. Membrane voltage fluctuations reduce spike frequency adaptation and preserve output gain in CA1 pyramidal neurons in a high-conductance state. J Neurosci 31: 3880–3893, 2011.

Francis JT, Gluckman BJ, Schiff SJ. Sensitivity of neurons to weak electric fields [Online]. J Neurosci 23: 7255–7261, 2003. http://www.jneurosci.org/content/23/19/7255.

Fröhlich F, McCormick DA. Endogenous Electric Fields May Guide Neocortical Network Activity. Neuron 67: 129–143, 2010.

Helfrich RF, Knepper H, Nolte G, Strüber D, Rach S, Herrmann CS, Schneider TR, Engel AK. Selective Modulation of Interhemispheric Functional Connectivity by HD-tACS Shapes Perception. PLoS Biol 12, 2014a.

Helfrich RF, Schneider TR, Rach S, Trautmann-Lengsfeld SA, Engel AK, Herrmann CS. Entrainment of Brain Oscillations by Transcranial Alternating Current Stimulation. Curr Biol 24: 333–339, 2014b.

Herrmann CS, Rach S, Neuling T, Strüber D. Transcranial alternating current stimulation: a review of the underlying mechanisms and modulation of cognitive processes. Front Hum Neurosci 7: 279, 2013.

Huang Y, Datta A, Bikson M, Parra LC. Realistic vOlumetric-Approach to Simulate Transcranial Electric Stimulation – ROAST – a fully automated open-source pipeline. J. Neural Eng. (May 9, 2019). doi:10.1088/1741-2552/ab208d.

Huang Y, Liu AA, Lafon B, Friedman D, Dayan M, Wang X, Bikson M, Doyle WK, Devinsky O, Parra LC. Measurements and models of electric fields in the in vivo human brain during transcranial electric stimulation. Elife 6: 1–27, 2017.

Huang Y, Parra LC, Haufe S. The New York Head—A precise standardized volume conductor model for EEG source localization and tES targeting. Neuroimage 140: 150–162, 2016.

Huk AC, Ress D, Heeger DJ. Neuronal Basis of the Motion Aftereffect Reconsidered. Neuron 32: 161–172, 2001.

Ito T, Kulkarni KR, Schultz DH, Mill RD, Chen RH, Solomyak LI, Cole MW. Cognitive task information is transferred between brain regions via resting-state network topology. Nat Commun 8: 1–13, 2017.

Kar K. Commentary: On the possible role of stimulation duration for after-effects of transcranial alternating current stimulation. Front Syst Neurosci 9: 311, 2015.

Kar K, Duijnhouwer J, Krekelberg B. Transcranial Alternating Current Stimulation Attenuates Neuronal Adaptation. J Neurosci 37: 2325–2335, 2017.

Kar K, Krekelberg B. Transcranial electrical stimulation over visual cortex evokes phosphenes with a retinal origin. J Neurophysiol 108: 2173–8, 2012.

Kar K, Krekelberg B. Transcranial alternating current stimulation attenuates visual motion adaptation. J Neurosci 34, 2014a.

Kar K, Krekelberg B. Transcranial alternating current stimulation attenuates visual motion adaptation. J Neurosci 34: 7334–40, 2014b.

Kar K, Krekelberg B. Testing the assumptions underlying fMRI adaptation using intracortical recordings in area MT. Cortex 80: 1–14, 2016.

Krause MR, Vieira PG, Csorba BA, Pilly PK, Pack CC. Transcranial alternating current stimulation entrains single-neuron activity in the primate brain. Proc. Natl. Acad. Sci. U. S. A. (2019). doi:10.1073/pnas.1815958116.

Krekelberg B, Boynton GM, van Wezel RJA. Adaptation: from single cells to BOLD signals. Trends Neurosci 29, 2006a.

Krekelberg B, van Wezel RJA, Albright TD. Adaptation in macaque MT reduces perceived speed and improves speed discrimination. J Neurophysiol 95: 255–70, 2006b.

Laakso I, Hirata A. Computational analysis shows why transcranial alternating current stimulation induces retinal phosphenes. J Neural Eng 10: 046009 (9pp), 2013.

Liu A, Greg S, Henin R, Krause M, Lucas C, Parra LC. Direct effects of transcranial electrical stimulation on the brain. Alexander Opitz Bart Krekelb Antal Berényi 1172, 2018.

Opitz A, Falchier A, Yan C, Yeagle E, Linn G. Spatiotemporal structure of intracranial electric fields induced by transcranial electric stimulation in human and nonhuman primates. Sci Rep 6: 1–11, 2016.

Ozen S, Sirota A, Belluscio MA, Anastassiou CA, Stark E, Koch C, Buzsáki G. Transcranial electric stimulation entrains cortical neuronal populations in rats. J Neurosci 30: 11476–85, 2010.

Patterson CA, Duijnhouwer J, Wissig SC, Krekelberg B, Kohn A. Similar adaptation effects in primary visual cortex and area MT of the macaque monkey under matched stimulus conditions. J Neurophysiol 111: 1203–13, 2014.

Power JD, Cohen AL, Nelson SM, Wig GS, Barnes KA, Church JA, Vogel AC, Laumann TO, Miezin FM, Schlaggar BL, Petersen SE. Functional Network Organization of the Human Brain. Neuron 72: 665–678, 2011.

Rezec A, Krekelberg B, Dobkins KR. Attention enhances adaptability: Evidence from motion adaptation experiments. Vision Res 44: 3035–3044, 2004.

Rubinov M, Sporns O. Complex network measures of brain connectivity: Uses and interpretations. Neuroimage 52: 1059–1069, 2010.

Scheeringa R, Petersson KM, Kleinschmidt A, Jensen O, Bastiaansen MCM. EEG lpha power modulation of fMRI restingstate connectivity. Brain Connect 2: 254–64, 2012.

Schutter DJLG, Hortensius R. Retinal origin of phosphenes to transcranial alternating current stimulation. Clin Neurophysiol 121: 1080–1084, 2010.

Stroud AC, LeDue EE, Crowder NA. Orientation specificity of contrast adaptation in mouse primary visual cortex. J Neurophysiol 108: 1381–1391, 2012.

Strüber D, Rach S, Neuling T, Herrmann CS, Kar K. On the possible role of stimulation duration for after-effects of transcranial alternating current stimulation. Front Cell Neurosci 9: 311, 2015.

Tootell RB, Reppas JB, Dale AM, Look RB, Sereno MI, Malach R, Brady TJ, Rosen BR. Visual motion aftereffect in human cortical area MT revealed by functional magnetic resonance imaging. Nature 375: 139–41, 1995.

Vosskuhl J, Huster RJ, Herrmann CS. BOLD signal effects of transcranial alternating current stimulation (tACS) in the alpha range: A concurrent tACS?fMRI study. Neuroimage 140: 118–125, 2016.

Zaehle T, Rach S, Herrmann CS. Transcranial Alternating Current Stimulation Enhances Individual Alpha Activity in Human EEG. PLoS One 5: 1–7, 2010.

